# Identification of *avaC* from infant gut microbial isolates that convert 5AVA to 2-piperidone with high efficiency

**DOI:** 10.1101/2023.11.16.567476

**Authors:** Qiudi Zhou, Lihui Feng

**Affiliations:** Institute of Pediatrics, Children’s Hospital of Fudan University, Institutes of Biomedical Sciences, Fudan University, Shanghai, 200032, China

**Keywords:** 2-piperidone, gut microbiota metabolism, cross-feeding

## Abstract

2-Piperidone has been identified as a biomarker for various human diseases, but its origin *in vivo* remains poorly understood. Furthermore, 2-piperidone is a crucial industrial raw material, and thus the discovery of novel efficient 2-piperidone synthases may have an important application in its biosynthesis. In this study, we aimed to identify the bacterial source and metabolic mechanism of 2-piperidone from a previously generated infant gut microbial culture collection. We found that 2-piperidone could be produced from 5AVA by four bacterial strains, including *Collinsella aerofaciens* LFYP39, *Collinsella intestinalis* LFYP54, *Clostridium bolteae* LFYP116, and *Clostridium hathewayi* LFYP18 from 51 bacterial strains. Furthermore, 2-piperidone could be synthesized from proline by cross-feeding between *Clostridium difficile* LFYP43 and *Collinsella intestinalis* LFYP54. We employed a gain-of-function library to identify the gene *avaC* (5-aminovaleric acid cyclase) which can catalyze 5AVA to produce 2-piperidone in *C. intestinalis* LFYP54. Homologous genes of *avaC* were also identified and validated in the other three bacterial strains. Gene *avaC* exhibits a broad distribution in the natural environmental bacteria. Overall, our research identified the gut bacterial strains and the genes that are responsible for the production of 2-piperidone. This study may facilitate the prediction of 2-piperidone-related disease risks through the analysis of gut bacterial community composition, and enhance the efficiency of 2-piperidone in its biosynthesis in industry.

**Importance:** In recent decades, it has become a consensus that gut microbiota can affect host health through producing metabolites. However, the complexity of gut flora composition makes finding the sources of the particular metabolite challenging. 2-piperidone is a biomarker for various diseases and a highly valued raw material for nylons. In this study, we identified the gut bacterial strains that can transform 5AVA to 2-piperidone. A novel 2-piperidone synthase gene, *avaC*, was also identified and characterized. These findings provided new insights into the potential connection between 2-piperidone related diseases and the intestinal flora, as well as a possible novel approach for more efficient biosynthesis of 2-piperidone in industry.

## Introduction

The gut microbiota utilizes nutrients ingested by the host to produce thousands of small molecule metabolites through diverse metabolic activities (1). The small molecule metabolites produced by the gut microbiota not only accumulate in the gastrointestinal tract but some could also enter the bloodstream and reach various organs of the host (2–4). Amino acids are essential nutrients for the human body. The gut microbiota can produce various small-molecule metabolites through amino acid metabolism to influence host physiology, such as polyamines (5, 6), indoles (7, 8), GABA (9), and so on. However, identifying the metabolic pathways of amino acid-derived small molecule metabolites is a challenge, and such effort may enable predicting the composition of the metabolites *in vivo* directly from sequencing data (10).

2-Piperidone is a lactam molecule that is cyclized from ω-amino acids. It was initially discovered as the product of lysine oxidation by L-amino acid oxidase in turkey liver, particularly in the absence of catalase (11). In addition, 2-piperidone has been detected in metabolite extracts from other animal body fluids (12–15). The biological activities of 2-piperidone include antifungal activity (16), insecticidal activity (17), and anticonvulsant effect in mammals (18). In a previous study using gnotobiotic mice inoculated with different gut microbial consortia (19), the amount of 2-piperidone in the cecum and serum of mice inoculated with certain microbial consortia was significantly higher than germ-free mice (unpublished data), suggesting that 2-piperidone could be produced by human gut microbial isolates. There is evidence that 2-piperidone is associated with human diseases, including epithelial ovarian cancer (EOC) (20, 21), Crohn’s disease (CD) (22–24), primary biliary cirrhosis (PBC) (25) and esophageal squamous cell carcinoma (ESCC) (26). Thus, investigating the metabolic pathway of 2-piperidone may enable prediction of disease risks through microbiome composition.

In addition, 2-piperidone is a fundamental scaffold for the synthesis of various chemical products, including high-value nylon-5 and nylon-6,5 (27–31). 2-piperidone is usually prepared from 5-aminovaleric acid (5AVA) as the substrate in biosynthetic pathways (32). The cyclization enzymes mainly used to produce 2-piperidone include ORF26, Act and CaiC, all of which come from bacteria (33, 34). ORF26, an enzyme encoded by *Streptomyces aizunensis*, can mediate the activation of 5AVA via ATP and facilitate the formation of 2-piperidone (33). Act, which belongs to the β-alanine CoA transferase family from *Clostridium propionicum*, can catalyze the activation of 5AVA using Acyl-CoA as a coenzyme and induce its subsequent cyclization to 2-piperidone (34). CaiC, a crotonobetaine CoA ligase, is encoded by *Escherichia coli* and is capable of catalyzing the cyclization of 5-aminovaleric acid into 2-piperidone. The cyclization step is rate-limiting in the biosynthesis of 2-piperidone (32). Exploring the metabolic pathway from the gut microbiota that produces 2-piperidone may reveal novel approaches for synthesizing 2-piperidone in industry.

To identify the metabolic pathway of 2-piperidone, we analyzed the culture supernatant of 51 intestinal bacteria strains isolated from an infant and identified four strains, including *Collinsella aerofaciens* LFYP39, *Collinsella intestinalis* LFYP54, *Clostridium bolteae* LFYP116, and *Clostridium hathewayi* LFYP18, which could produce 2-piperidone from 5AVA. In addition, we established an *in vitro* cross-feeding system, through which *Clostridium difficile* LFYP43 and *Collinsella intestinalis* LFYP54 produced 2-piperidone using proline as the substrate. Subsequently, we identified *avaC* as the gene that was able to convert 5AVA to 2-piperidone from the above four strains by genomic DNA library screening and bioinformatic analysis. The determination of the metabolic pathway of 2-piperidone from the gut microbiota helps to understand the correlation between gut microbiota and 2-piperidone related diseases, as well as providing a new approach to the biosynthesis of 2-piperidone in industry.

## Results

### Four gut microbial species produce 2-piperidone from 5AVA

Based on the molecular structure of 2-piperidone and previous studies, we hypothesized that 5-aminovaleric acid (5AVA) could be a precursor of 2-piperidone *in vivo*. To identify which gut bacterial strains could metabolize 5AVA to produce 2-piperidone in our infant gut bacterial culture collection, we chose 51 gut bacterial strains representing five major gut microbiota phylum, incubated each of them with or without 5AVA *in vitro* and measured the amount of 2-piperidone in the supernatant by LC-MS/MS (see Methods). We found that four bacterial species, including *Collinsella intestinalis* LFYP54, *Clostridium bolteae* LFYP116, *Collinsella aerofaciens* LFYP39 and *Clostridium hathewayi* LFYP18, could produce 2-piperidone when incubated with 5AVA (Fig.1A). Phylogenetic analysis of the 51 intestinal bacterial strains based on the sequence of 16S ribosomal RNA genes revealed that the two strains of *Collinsella*, *C. intestinalis* LFYP54 and *C. aerofaciens* LFYP39, were located on the same branch of the evolutionary tree, as were the two strains of *Clostridium*, *C. bolteae* LFYP116 and *C. hathewayi* LFYP18. However, the relationship between *Collinsella* and *Clostridium* is not adjacent (Fig. 1A).

**Figure 1.**
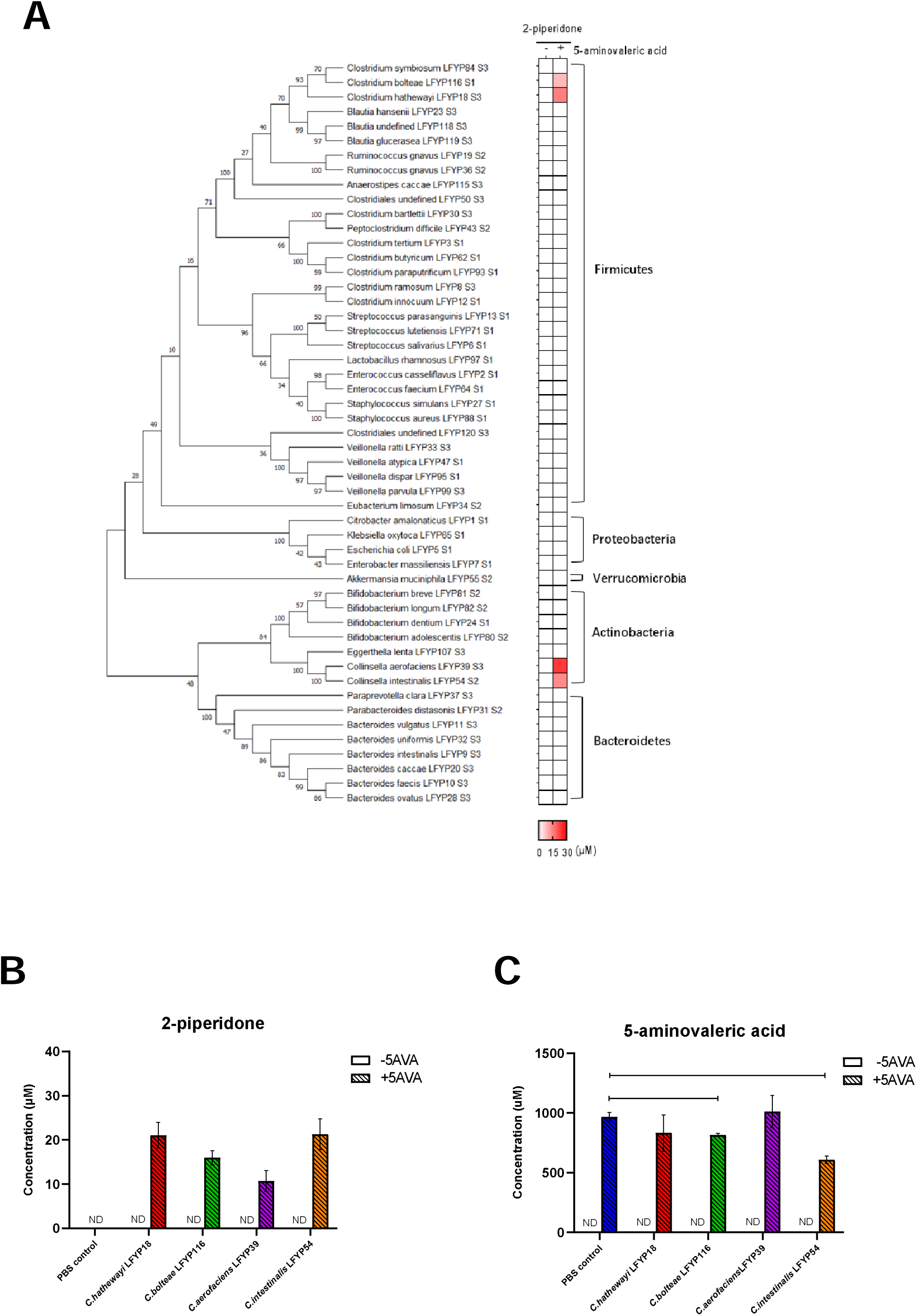
Four gut microbial species convert 5AVA to 2-piperidone. **A.** Heat map showing the mean concentration of 2-piperidone in the supernatant from 51 gut bacterial strains incubated without or with 5AVA (n=3 biological replicates). The phylogenetic tree of the 51 gut bacterial strains was constructed by Maximum Likelihood method and Tamura-Nei model based on 16S rDNA sequences. The percentage of trees in which the associated taxa clustered together is shown next to the branches. **B.** Concentration of 2-piperidone in the supernatant of PBS control and the four gut microbial strains incubated without or with 1 mM 5AVA. **C.** Concentration of 5AVA in the supernatant of PBS control and the four gut microbial strains incubated without or with 1 mM 5AVA. **B, C** Error bars represent the standard error of mean from three biological replicates. ND, not detected. *, p<0.05 and **, p<0.01 by unpaired t-test.

To verify the above positive results, we performed additional biological replicates. The results showed that *C. intestinalis* LFYP54, *C. bolteae* LFYP116, *C. aerofaciens* LFYP39 or *C. hathewayi* LFYP18 produced 20 μM, 17 μM, 10 μM and 20 μM 2-piperidone, respectively, after co-incubated with 5AVA (Fig. 1B). The conversion rates of 5AVA to 2-piperidone were comparable across the four strains, at 2%, 1.7%, 1%, and 2%, respectively. Only *C. intestinalis* LFYP54 and *C. bolteae* LFYP116 showed obvious consumption of 5AVA (Fig. 1C), which might be involved in other metabolic activities.

### *C. difficile* and *C. intestinalis* collaborate to produce 2-piperidone from proline *in vitro*

The above results showed that four infant gut microbial strains could produce 2-piperidone from 5AVA. Next, we wondered about the source of 5AVA. Previous studies have shown that *Clostridium difficile* and *Clostridium sporogenes* could produce 5AVA from proline (3,4). We hypothesized that *C. difficile* LFYP43 from our gut bacterial culture collection could also produce 5AVA using proline as the substrate. Using similar *in vitro* system as described above, we incubated *C. difficile* LFYP43 with proline. The results showed that *C. difficile* LFYP43 produced 500 μM 5AVA after incubated with 1000 μM proline (Fig. 2A). As a control, *C. difficile* LFYP43 incubated without proline produced 20 μM 5AVA (Fig. 2A). Meanwhile, the amount of proline decreased by almost 50% when incubated with *C. difficile* LFYP43 (Fig. 2B).

**Figure 2.**
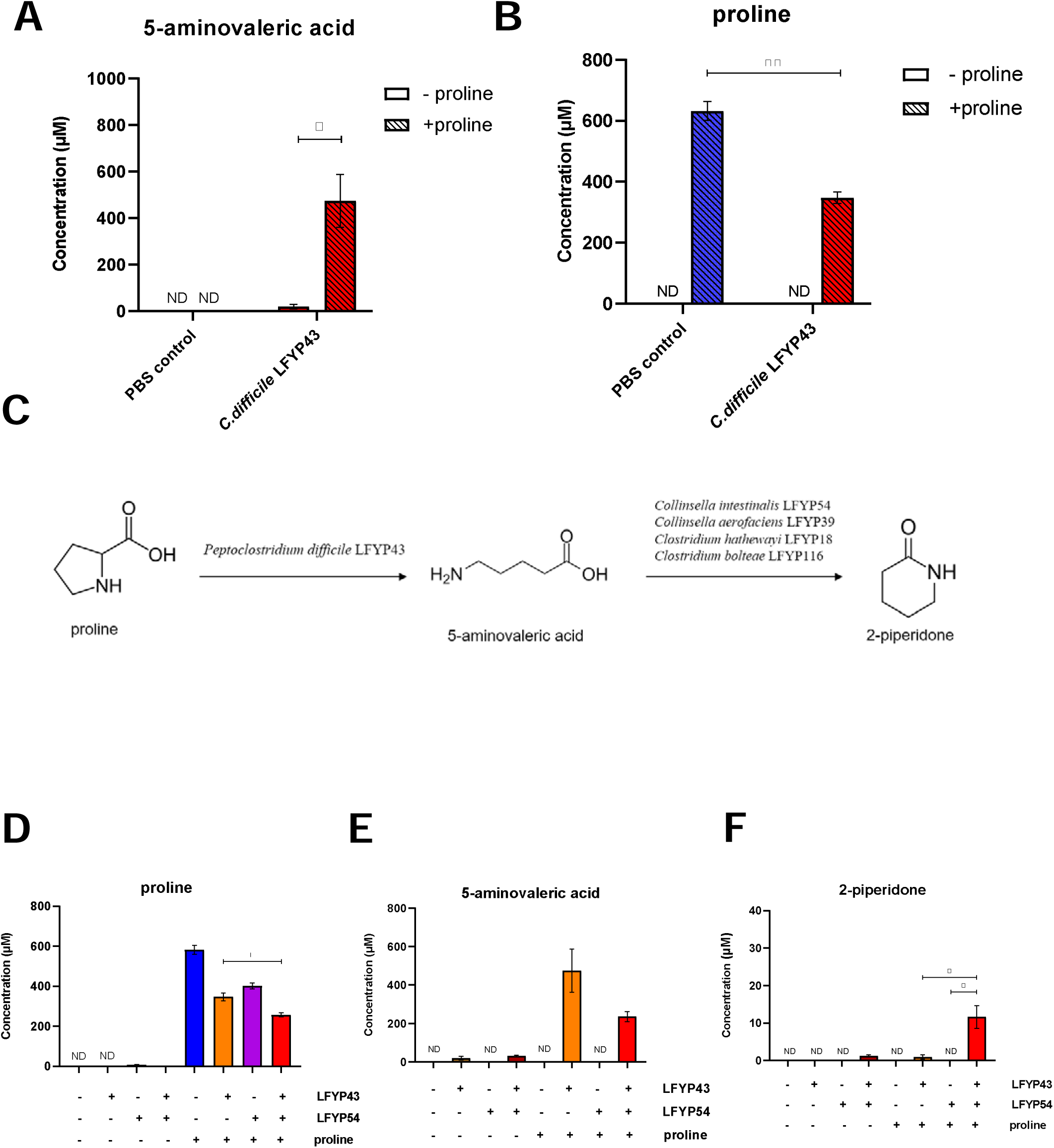
Gut Microbial strains produce 2-piperidone from proline through cross-feeding *in vitro*. Concentration of 5AVA (**A**) and proline (**B**) in the supernatant of PBS control and *C. difficile* LFYP43 incubated without or with 1 mM proline. **C.** A proposed cross-feeding pathway between *C. difficile* LFYP43 and the four 2-piperidone-producing strains. Concentration of proline (**D**), 5AVA (**E**) and 2-piperidone (**F**) in the supernatant of *C. difficile* LFYP43 and *C. intestinalis* LFYP54 under different incubation conditions. Error bars represent the standard error of mean from three biological replicates. ND, not detected. *, p<0.05 and **, p<0.01 by unpaired t-test.

Based on the results above, we hypothesized that metabolic cross-feeding could occur between *C. difficile* LFYP43 and one of the four bacterial species that could produce 2-piperidone from 5AVA. Specifically, when co-incubated, proline could be metabolized to 5AVA by *C. difficile* LFYP43, and 5AVA could further be converted to 2-piperidone by *C. intestinalis* LFYP54, *C. bolteae* LFYP116, *C. aerofaciens* LFYP39 or *C. hathewayi* LFYP18 (Fig. 2C). To test this hypothesis, similar incubation experiments were performed with both *C. difficile* LFYP43 and *C. intestinalis* LFYP54 with the addition of proline as the substrate *in vitro*. LC-MS analysis of the supernatant showed that consumption of proline was 56% when *C. intestinalis* LFYP54 and *C. difficile* LFYP43 were co-incubated with 1000 μM proline, whereas *C. difficile* LFYP43 alone resulted in 40% consumption of proline (Fig. 2D). Meanwhile, when *C. difficile* LFYP43 and *C. intestinalis* LFYP54 were co-incubated with 1000 μ M proline, the concentration of 5AVA was lower (236 μM) compared to incubation with *C. difficile* LFYP43 alone (475 μM), presumably due to the further metabolization of 5AVA by *C. intestinalis* LFYP54 (Fig. 2E). Incubation of proline and *C. intestinalis* LFYP54 or *C. difficile* LFYP43 alone almost did not produce 2-piperidone. However, when *C. difficile* LFYP43 and *C. intestinalis* LFYP54 were co-incubated with 1000 μM proline, they produced 12 μM 2-piperidone (Fig. 2F). These results supported that cross-feeding interactions existed in *C. difficile* LFYP43 and *C. intestinalis* LFYP54, in that these two strains cooperated to produce 2-piperidone using proline as the substrate.

### Gene *avaC* from *C. intestinalis* LFYP54 converts 5AVA to 2-piperidone

The above study revealed four bacterial strains that can convert 5AVA to 2-piperidone, which has not been reported before. Next, we wanted to identify the gene that is responsible for catalyzing this reaction. It was reported that ORF26 in *Streptomyces aizunensis*, β-alanine CoA transferase (Act) in *Clostridium propionicum* and carnitine CoA ligase (CaiC) in *Escherichia coli* could cyclize 5AVA to form 2-piperidone [7,8]. We aligned the sequences of *orf26*, *act* and *caiC* with the whole-genome sequencing data of the four bacterial strains. No homologous sequence were found, suggesting that these four newly identified bacterial strains use a different mechanism.

To identify the gene product that could produce 2-piperidone from 5AVA in the four bacterial strains, we employed a previously described two-plasmid biosensor system [10]. The two-plasmid biosensor system in *E. coli* contained a biosensor plasmid (pLacSens) and a production plasmid. The pLacSens expresses green fluorescent protein (GFP) upon sensing 2-piperidone by the transcription factor OplR. The production plasmid was inserted with random DNA sequences under the control of the PLlacO1 promoter. If the DNA sequence inserted in the production plasmid could produce 2-piperidone from 5AVA, green fluorescence would be detected (Fig. 3A). To evaluate the sensitivity of the biosensor plasmid for detecting 2-piperidone, different concentrations of 2-piperidone were added to the culture medium of *E. coli* harboring the pLacSens plasmid and GFP was measured. The results showed that GFP was ∼2 fold when 1 μM of 2-piperidone was added compared to the control group without adding 2-piperidone. Moreover, GFP reached saturation when 200 μM of 2-piperidone was added, which is ∼6 fold of the control group (Fig. S1, squares). Next, two control strains were constructed. The negative control strain carries pLacSens and an empty production plasmid, whereas the positive control carries pLacSens and a production plasmid inserted with *orf26*. 2-piperidone can also induce GFP production in *E. coli* carrying the two-plasmid system (Fig. S1). To evaluate the effectiveness of the system, the negative and positive control strains were cultured in LB medium with and without 5AVA, and GFP and 2-piperidone levels were measured (Fig. S3). In the positive control strain, GFP signal increased ∼1.4 fold with 5AVA compared to without 5AVA; whereas in the negative control strain, GFP signal increased ∼0.7 fold (Fig. S2A). LC-MS/MS revealed that the positive control strain produced ∼14 μM 2-piperidone after incubating with 5 mM 5AVA and the negative control strain did not produce detectable levels of 2-piperidone (Fig. S2B). These results provided the basis for the availability of the two-plasmid screening system.

**Figure 3.**
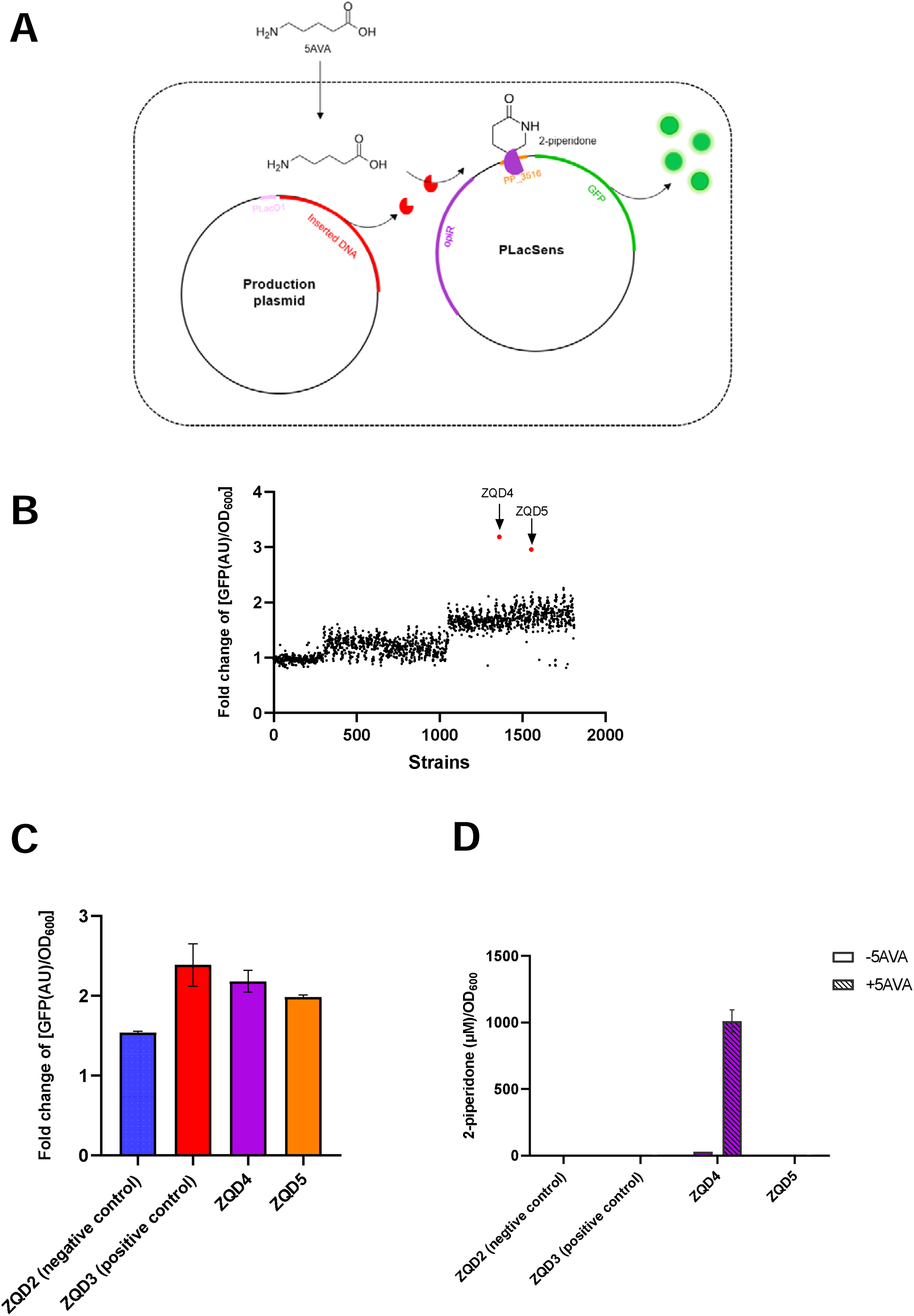
Identification of the genomic sequence in *C. intestinalis* LFYP54 which converts 5AVA to 2-piperidone. **A.** Schematic plot illustrating the principle of the two-plasmid biosensor screening system. **B.** Fold-change of [GFP(AU)/OD_600_] values (see Methods for definition) from *E. coli* single colonies. Colonies with significant higher fold-change values than the mean were colored red (ZQD4 and ZQD5). **C.** Fold-change of [GFP(AU)/OD_600_] values from the negative and positive control strains, ZQD4 and ZQD5. **D.** Concentration of 2-piperidone per OD_600_ cell in the supernatant when strains were cultured without or with 5 mM 5AVA. Error bars represent the standard error of mean from three biological replicates.

Next, since the efficiency of transforming 5AVA to 2-piperidone is similar between the four bacterial strains, one of them, *C. intestinalis* LFYP54, was selected as the target gene screening object. Genomic DNA fragments of *C. intestinalis* LFYP54 were inserted to the empty production plasmid (See Methods). ∼1800 colonies grown on a plate containing antibiotics were picked. Colonies were cultured in the presence and absence of 5AVA, and the GFP signal was measured for each colony culture. Fold-change of GFP were calculated for each colony (See Methods). Among the1800 colonies, two strains, named ZQD4 and ZQD5, showed significantly higher (∼3 fold) fold-change of GFP values compared to the other colonies (Fig. 3B). ZQD4 and ZQD5 were repurified and the replication experiments confirmed the screening results (Fig. 3C). LC-MS/MS analysis was also performed to confirm the production of 2-piperidone from ZQD4 and ZQD5, and surprisingly, only ZQD4 produced 2-piperidone (Fig. 3D). It was worth mentioning that the amount of 2-piperidone produced by ZQD4 is 1 mM after incubating with 5 mM of 5AVA, which was 50-fold of that of the positive control strain (0.002 mM), indicating that the gene carried in ZQD4 had a much greater ability to convert 5AVA to 2-piperidone than the previously reported gene *orf26*.

The inserted DNA sequence in the production plasmid of ZQD4 was sequenced and aligned with the whole-genome sequence of *C. intestinalis* LFYP54. The results showed that the 2931bp inserted DNA fragment mainly included two genes, *CILFYP54_00697* and incomplete *ypdA* (Fig. 4A). *CILFYP54_00697* encodes a hypothetical protein annotated as an amidohydrolase family protein and *ypdA* encodes a sensor histidine kinase (YpdA). To identify which gene could convert 5AVA to 2-piperidone, we cloned each of them to the production plasmid. Heterologous expression of *CILFYP54_00697* alone resulted in a 3-fold increase of GFP with 5AVA compared to without 5AVA, similar to that of expressing the 2931bp sequence or *CILFYP54_00697* and y*pdA* together, whereas expressing the complete *ypdA* gene resulted in ∼1.3-fold change in GFP (Fig. 4B). LC-MS/MS analysis revealed that expressing *CILFYP54_00697* produced 2.2 mM 2-piperidone in the supernatant when 5 mM 5AVA was added in the medium, whereas expressing *ypdA* alone did not produce2-piperidone (Fig. 4C). Therefore, gene *CILFYP54_00697* is capable of catalyzing the dehydration and cyclization of 5AVA to produce 2-piperidone. We named gene *CILFYP54_00697* as *avaC* (5-aminovaleric acid cyclase). Notably, co-expression of *avaC* and *ypdA* resulted in the production of 3.5 mM 2-piperidone, which was significantly higher than expressing *avaC* alone (Fig. 4C), suggesting that the presence of *ypdA* may promote the ability of *avaC* to convert 5AVA to 2-piperidone.

**Figure 4.**
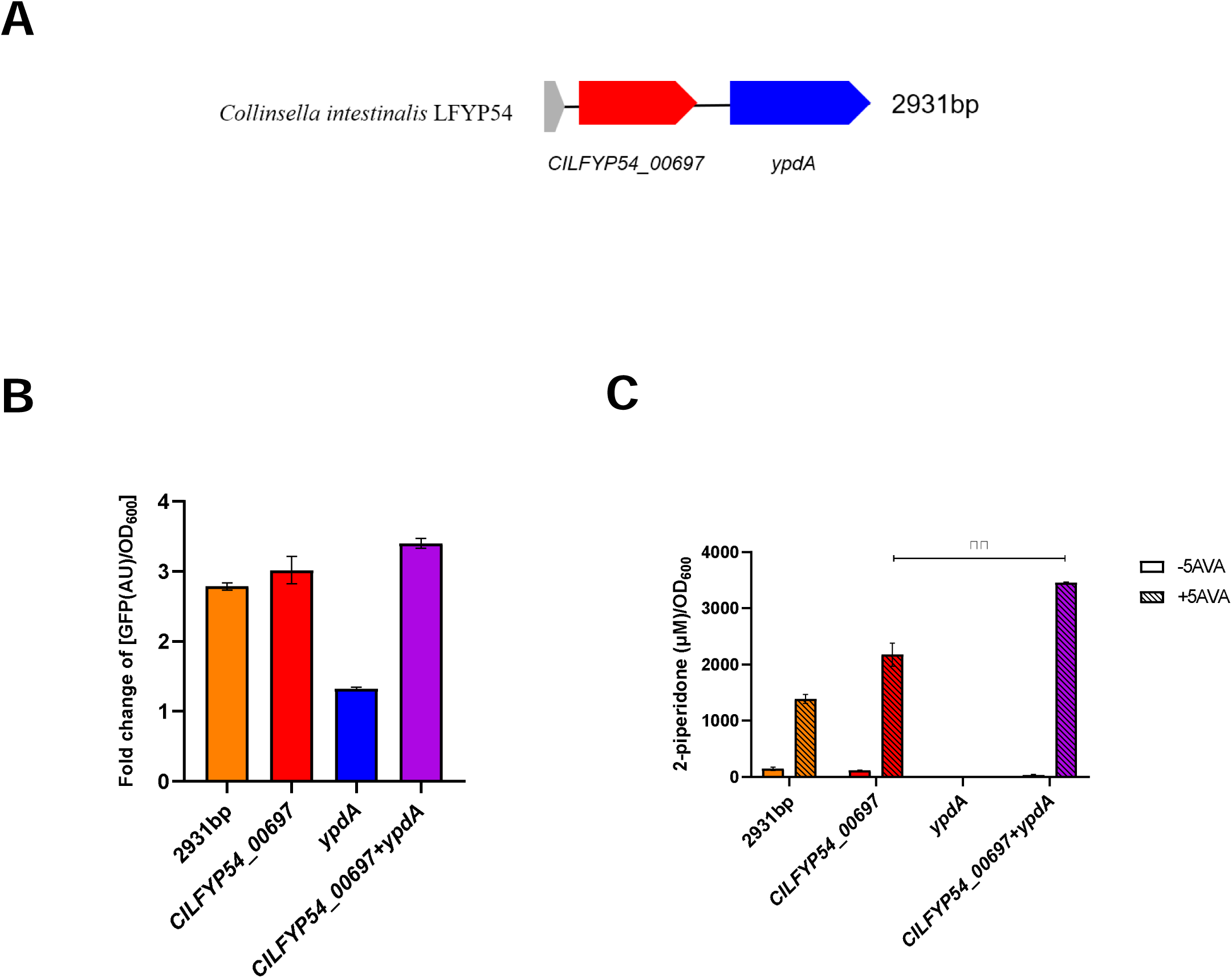
Gene *avaC* in *C. intestinalla* LFYP54 converts 5AVA to 2-piperidone. **A.** Schematic plot showing the 2931 bp DNA sequence inserted into the production plasmid in strain ZQD4. *CILFYP54_00697* encodes a hypothetical protein and *ypdA* encodes the sensor histidine kinase YpdA. **B.** Fold change of [GFP(AU)/OD_600_] of *E. coli* strains heterologously expressing the different DNA sequences as indicated. **C.** Concentration of 2-piperidone per OD_600_ cell in the supernatant when strains were cultured without or with 5 mM 5AVA. Error bars represent the standard error of mean from three biological replicates. **, p<0.01 by unpaired t-test.

### Additional genes in gut microbial strains that convert 5AVA to 2-piperidone

We have identified the gene *avaC* that could convert 5AVA to 2-piperidone in *C. intestinalis* LFYP54. Next, we intended to characterize other genes with the same function as *avaC* in *C. aerofaciens* LFYP39, *C. bolteae* LFYP116 and *C. hathewayi* LFYP18. The nucleotide sequence of *avaC* from *C. intestinalis* LFYP54 was aligned with whole-genome sequences of the other three strains. *C. aerofaciens* LFYP39, which is phylogenetically related to LFYP54, contains a gene (*CALFYP39_00071*) that shares highly similar sequences with *avaC*. However, *C. hathewayi* LFYP18 and *C. bolteae* LFYP116, which are distantly related to *C. intestinalis* LFYP54, harbor genes (*CHLFYP18_04700* and *CBLFYP116_02895*, respectively) that exhibit more dissimilar sequences (Fig 5A). The protein sequence alignment results showed that the three protein sequences are highly similar to the AvaC protein. These proteins were annotated as a set of enzymes that catalyze the hydrolysis of a wide range of substrates bearing amide or ester functional groups at carbon [9], which is the reverse of the cyclization reaction.

**Figure 5.**
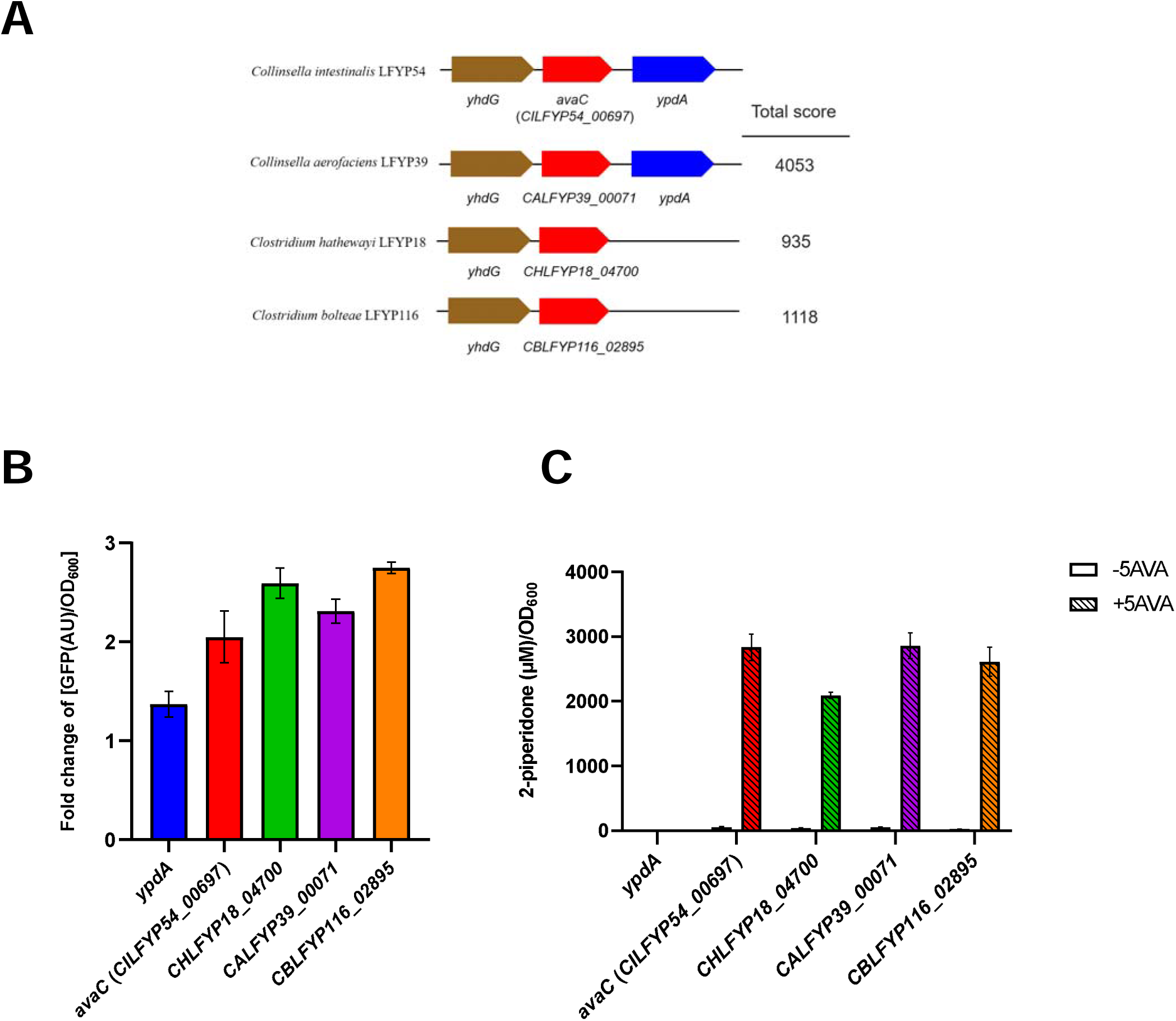
Homologous genes of *avaC* in other intestinal bacteria. **A.** Similar gene clusters containing gene *avaC* from other intestinal bacteria. *yhdG* encodes putative amino acid permease. *avaC* (*CILFYP54_00697*) encodes 5-aminovaleric acid cyclase. Similarity score was calculated by Nucleotide Basic Local Alignment Search Tool (BLAST). **B.** Fold change of [GFP(AU)/OD_600_] of *E. coli* strains with the two-plasmid biosensor system expressing the homologous genes as indicated. **C.** Concentration of 2-piperidone per OD_600_ cell in the supernatant of *E. coli* strains expressing the homologous genes incubated without or with 5 mM 5AVA.

To test the function of these additional genes, we cloned them to the production plasmid. The *ypdA* gene was employed as a negative control. As expected, the other three genes resulted in similar GFP signal and 2-piperidone production to *avaC* from *C. intestinalis* LFYP54 (Fig. 5B, C). We named *CALFYP39_00071*, *CHLFYP18_04700* and *CBLFYP116_02895* as *avaC* as well.

To further know the distribution of this type of enzyme in bacteria, we performed a sequence similarity search using the four a*vaC* genes in the National Center for Biotechnology Information (NCBI) GenBank database (Table S4). The results showed that: (1) Their homologous genes were largely overlapping, with varying scores of sequence similarity among them; (2) The homologous genes were distributed in 5 phylum and mainly in *Bacillota* and *Actinomycetota;* (3) The bacteria containing homologous genes are widely distributed throughout various environments, with the majority belonging to gastrointestinal tract bacteria and some existing in natural or engineered environments.

## Discussion

In this study, we identified four intestinal bacterial strains, *C. intestinalis* LFYP54, *C. bolteae* LFYP116, *C. aerofaciens* LFYP39, and *C. hathewayi* LFYP18, which could produce 2-piperidone from 5AVA. In addition, we showed that 2-piperidone could be produced from proline by cross-feeding between *C. difficile* LFYP43 and *C. intestinalis* LFYP54. Furthermore, a new gene, *avaC*, in *C. intestinalis* LFYP54, *C. bolteae* LFYP116, *C. aerofaciens* LFYP39, and *C. hathewayi* LFYP18 was identified as having the function of converting 5AVA to 2-piperidone. Bioinformatic analysis revealed a broad distribution of *avaC* in the natural environmental bacterial species.

2-piperidone has been identified as a biomarker for various diseases, such as epithelial ovarian cancer (EOC), inflammatory bowel disease (IBD), esophageal squamous cell carcinoma (ESCC), and others (20, 22, 26). However, the underlying mechanisms, especially its sources *in vivo* are largely unclear. Our study revealed that 2-piperidone could be produced by intestinal bacterial isolates. In particular, *C. intestinalis* LFYP54, *C. bolteae* LFYP116, *C. aerofaciens* LFYP39, and *C. hathewayi* LFYP18 could produce 2-piperidone using 5AVA as substrate. Furthermore, *C. difficile* LFYP43 and *C. intestinalis* LFYP54 could corporately produce 2-piperidone using proline as substrate. Similar bacterial species may have the same functions as the bacterial strains used in this study. Future studies to investigate the distribution of 2-piperidone in host organs and its functions *in vivo* would facilitate the understanding of the relationship between 2-piperidone and the related diseases. It may also be possible to determine the risk of related diseases by sequencing the composition of the gut microbiota.

2-piperidone is a precursor for the production of nylon-5,6 (35), and its biosynthesis approach has garnered high-profile attention. The development of efficient biocatalytic method for synthesizing 2-piperidone can facilitate the green production of 2-piperidone. To data, expression of different 5AVA cyclization enzymes, including Act, ORF26 and CaiC, in genetically modified strains under identical culture conditions resulted in similar 2-piperidone yields (32). Compared to *orf*26, the newly discovered gene, *avaC*, exhibits a 50-fold increase in yield in the cyclization of 5AVA to 2-piperidone in this study. The cyclization step of 5AVA is rate-limiting to the yield of 2-piperidone (32). Our data suggested that *avaC* may be used to break the bottleneck of biological synthesis of 2-piperidone. Furthermore, Act, ORF26 and CaiC are capable of cyclizing 4ABA and 6ACA to form four- and six-carbon lactams (36). These reactions are all intramolecular dehydration cyclization reactions of ω-amino acids. Four- and six-carbon lactams are also of significant industrial value. Whether avaC can cyclize other ω-amino acids remains to be explored.

## Materials and Methods

### Reagents used for this research

All of the reagents used in this work are listed in Table S1.

### Bacterial strains and culture condition

All of the strains used in this work are listed in Table S2.

51 gut bacterial strains previously derived from a healthy infant (19) were stocked at -80℃ in a 2 ml glass crimp top vial containing cell freezing medium (30% glycerol, 7% 10*PBS, 0.5 g/L cysteine-HCl) until use. The 51 strains were cultured in LYHBHI broth (37 g/L BHI, 5 g/L yeast extract, 1 g/L cellobiose, 1 g/L maltose, 0.5 g/L L-cysteine-HCl, 5 mg/L hemin) or LYHBHI broth supplemented with 0.1% (w/v) soluble starch and 0.5% (w/v) partially purified porcine stomach mucin at 37℃ in an anaerobic chamber from Coy Laboratory Products. The gas composition used in the anaerobic chamber is 75% nitrogen, 20% carbon dioxide and 5% hydrogen, and the hydrogen concentration in the chamber was maintained at ∼2%. For the experiment of precursor metabolite incubations, the medium was sub-cultured from an early stationary phase culture at an initial OD_600_ of ∼0.001.

All *E. coli* used in this work were cultured aerobically at 37 °C in LB (Luria-Bertani) broth or agar supplemented with kanamycin (100 mg/L), ampicillin (100 mg/L) or both of them as required overnight. Then the overnight culture was transferred into the fresh LB with antibiotic(s) as required at a starting OD_600_ of ∼0.01 OD_600_ was measured by GENESYS 30 spectrophotometer (Thermo Fisher).

### Plasmid construction and cloning

Plasmids, synthetic DNA sequences, and primers are listed in Table S3. Plasmids were isolated using the Plasmid DNA Mini Kit I (Omega). SnapGene was used to design the primers, which were ordered from Sangon Biotech. All DNA fragments used to construct the plasmid were chemically synthesized by Genewiz. PCR reactions were performed using high-fidelity DNA polymerase (AB clonal). Plasmids were constructed by connecting the different fragments using the One Step Cloning Kit (Novoprotein) and chemically transformed to *E. coli* DH10B.

The biosensor plasmid (pLacsens) was constructed by ligating the PCR linearized pEVS143 vector (prime 1,2) and the synthetic gene fragment 1 (Table S3). The positive control plasmid was constructed by ligating the synthetic DNA fragment 2 (Table S3) to the PCR-linearized pZE12 vector (prime 3,4). The negative control plasmid (pZE12 empty vector) was generated by ligating two PCR-linearized pZE12 plasmid vectors together (prime 5,6 and primer 7,8).

Construction of production plasmids. The production plasmid library was generated using a previously described method (37). Briefly, crude genomic DNA was extracted from *C. intestinalis* LFYP54 grown to early stationary phase in LYHBHI. Crude genomic DNA was treated with RNase A (Takara) to remove RNA and purified using the PCR Purification Kit (QIAquick). The pure genomic DNA was sheared by ultrasound and 2-8 kb fragments were extracted from a 0.7% agarose gel using the Gel Extraction Kit (Qiagen). The gel-recovered fragments were ligated to the linearized PZE12 vector (prime 3,4) by blunt-end ligation enzyme (Epicentre FastLinkTM kit) and ligation products in the range of 4-10 kb were extracted from a 0.7% agarose gel.

### Targeted gene screening assay

The recovered ligation products were transformed into *E. coli* with pLacSens electrocompetent cells. Single clones grown on LB agar plates were picked to a 384-well plate containing LB medium with Kanamycin and Ampicillin. Overnight cultures from the 384-well plate were diluted at 1:100 (v/v) to 96-well plates containing LB medium without or with 5AVA and incubated for 24 h. GFP and OD_600_ values were measured using the Synergy LX Multimode Reader (BioTek). Fold change of GFP/OD_600_, defined as 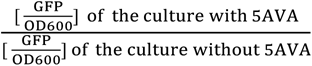, for the original clonal subculture in LB medium was determined. Strains with the highest fold change values of GFP/OD_600_ was selected for further LC-MS/MS analysis.

### Precursor metabolite incubation with resting cell suspensions

Precursor metabolite incubation experiments were performed by a previously described protocol (10). All operations were performed in the anaerobic chamber. In brief, overnight cultures of each gut bacterial strain were transferred at OD_600_ of ∼0.001 to fresh medium and grown to the early stationary phase. Then the cell pellet equivalent to 20 ml of OD_600_=1.0 from each strain for co-culture strains was harvested by centrifugation at 4500 rpm for 10 min. The cell pellet was washed twice with PBS, followed by resuspension in 4 ml of PBS. The cell resuspension was incubated at 37°C for 30 min to consume nutrition from the rich medium. The resuspension was divided equally into two portions and transferred to versatility tubes, with one portion receiving 1 mM (final concentration) metabolic precursor and the other untreated as negative control. Finally, the supernatant of the bacterial incubation was collected by centrifugation at 13300 rpm for 10 min and analyzed for metabolites using LC-MS/MS. To *C. bolteae* LFYP116, the strain supernatant was separated from the strain precipitate by filtering using 0.22 μm filter (Millipore).

### Liquid Chromatography-Tandem Mass Spectrometry (LC-MS/MS) quantitative analysis

All samples were centrifuged at 13300 rpm for 20 min before detection by LC-MS/MS. A Shimadzu LC with a Nucleodur C18 ISIS column (2× 250 mm) coupled to an AB SCIEX 4000 linear ion trap mass spectrometer was used to separate and detect the target compounds in samples. Two microliters of each sample were injected into the mobile phase via autosampler and separated at a flow rate of 0.2 ml*min^−1^. The mobile phase consisted of water with 0.1% formic acid (v/v) as eluent A and acetonitrile as eluent B. The gradient elution started with 100% eluent A for 4 min, followed by a linear gradient from 0 to 90%(v/v) eluent B in 7 min, and equilibrated with 90% eluent B for 2 min. Mass spectrometry was performed in positive ion mode under scan mode from 100 m/z to 400 m/z (For detailed parameters see Table S5). MRM parameters and retention times of the target compounds are shown in Table S6. The target compounds were quantified from calibration curves plotted with external standards. Some samples were analyzed by AB SCIEX 6500 triple-quadrupole liquid chromatography-mass spectrometry system.

### 16S rDNA phylogenetic tree construction

16S rRNA gene sequence (1249-1310 bp) was obtained by Sanger sequencing. The 16S rRNA gene sequences of 51 gut bacteria were aligned using the SILVA Aligner service (38). The aligned sequences were imported into MEGA X v10.2.6 software (39) to construct phylogenetic trees.

Evolutionary analysis by Maximum Likelihood method. The evolutionary history was inferred by using the Maximum Likelihood method and Tamura-Nei model (40). The tree with the highest log likelihood (-23769.34) is shown. The percentage of trees in which the associated taxa clustered together is shown next to the branches. Initial tree(s) for the heuristic search were obtained automatically by applying Neighbor-Join and BioNJ algorithms to a matrix of pairwise distances estimated using the Tamura-Nei model and then selecting the topology with superior log likelihood value. The tree is drawn to scale, with branch lengths measured in the number of substitutions per site. This analysis involved 51 nucleotide sequences. There were a total of 1434 positions in the final dataset. Evolutionary analyses were conducted in MEGA X (39).

### Homologous gene alignment

The Nucleotide Basic Local Alignment Search Tool (BLAST) was used to search the homologous genes of *CILFYP54_00697* or assess the homology of the gene cluster (*CILFYP54_00696* to *CILFYP54_00698*) in the genomes of *C. bolteae* LFYP116, *C. aerofaciens* LFYP39 and *C. hathewayi* LFYP18.

The Protein Basic Local Alignment Search Tool (BLAST) was used to search the NCBI GenBank database with the amino acid sequences of AvaC protein from *C. intestinalis* LFYP54, *C. aerofaciens* LFYP39, *C. hathewayi* LFYP18 and *C. bolteae* LFYP116 as queries. The protein sequences of identity >50% and coverage >80% were filtered in the search results, and the protein information was listed in Table S4.

## Acknowledgements

We thank members from the Feng lab for insightful discussions and the Mass Spectrometry Center of Institutes of Biomedical Sciences from Fudan University for technical supports. This work was supported by Zhuoshi Grant from Fudan University, Thousand Young Talents Program of China, Science and Technology Innovation Program of Shanghai grant 21ZR1480100.

We declare no conflicts of interest.

---

**Supplementary Figure 1.**
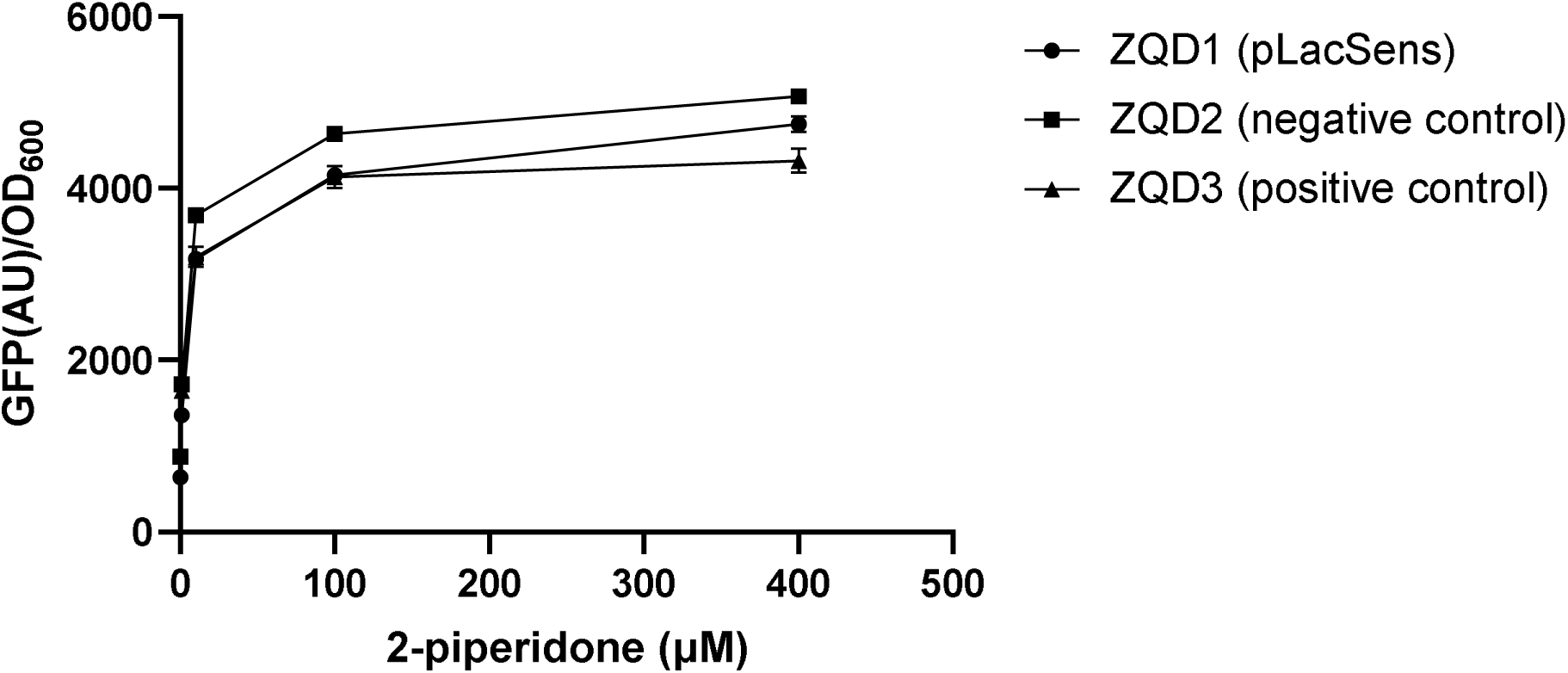
Sensitivity of the two-plasmid biosensor system to 2-piperidone. GFP(AU)/OD_600_ values of *E. coli* DH10B carrying different plasmids in response to various concentrations of 2-piperidone. (ZQD1: pLacSens; ZQD2 (negative control): pLacSens and an empty production plasmid; ZQD3 (positive control): pLacSens and a production plasmid with *orf26*. Error bars represent the standard error of mean from three biological replicates.

**Supplementary Figure 2.**
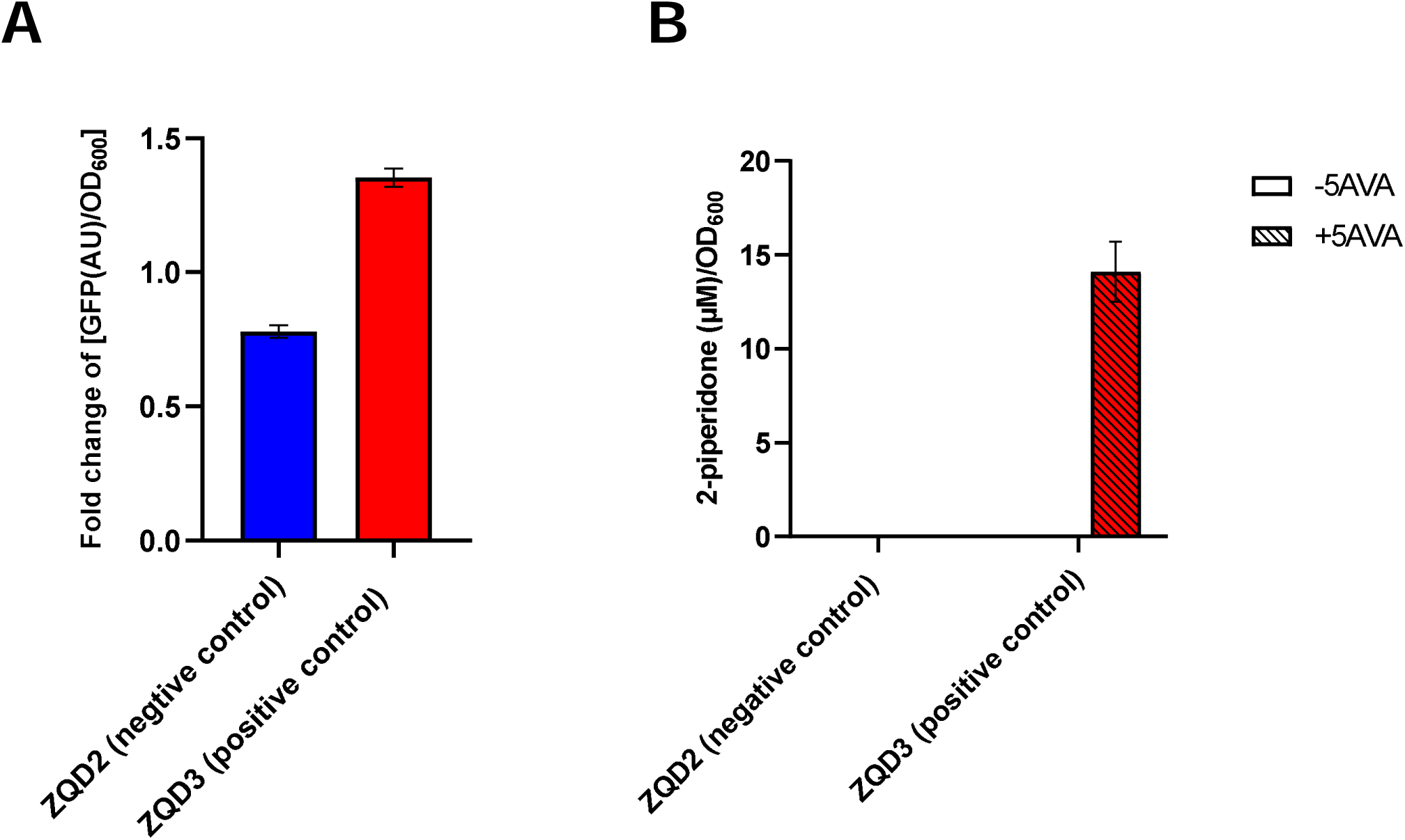
Control groups for the two-plasmid screening system. **A.** Fold change of [GFP(AU)/OD_600_] values of the positive control and negative control strains. **B.** Concentration of 2-piperidone per OD_600_ cell in the supernatant of the positive control and negative control strains incubated without or with 5 mM 5AVA. Error bars represent the standard error of mean from three biological replicates.

